# Learn to hear your prey: The role of associative learning on web building and hunting behaviour in black widow spiders (*Latrodectus hesperus*)

**DOI:** 10.1101/2025.07.04.663247

**Authors:** Stéphanie Moreau, Sarah Saneeibajgiran, Léo Korst, Pierre-Olivier Montiglio

## Abstract

Individuals from many populations vary consistently in their diet or behaviour. Associative learning could lead to such specialisation by improving the ability of predators to detect or recognize prey types or optimizing the sensory cues or spatial locations that they attend to. In this study, we assess the ability of Western black widow spiders for associative learning in response to vibrational prey cues. Black widow spiders are sedentary predators that detect, identify, and choose whether to attack preys caught in the web using the frequency and strength of vibrations. We conducted two experiments assessing the ability of individuals to associate specific frequencies or spatial locations on the web to the presence of a prey. We analyzed changes in web structure and attack behavior through learning. We hypothesized that individuals would adjust the structure of their webs and their responsiveness towards these frequencies in response to these associations. Spiders did not adjust the structure of their web nor their response to specific web locations when we applied prey items and vibrations at specific locations on the web. Instead, all spiders increased their responsiveness to vibrations at 250 Hz irrespective of their experimental treatment, but not towards 25Hz cues. Unexpectedly, exposing spiders to prey items associated with a 250Hz increased the effect of trap threads on responsiveness itself. Hence, repeated exposures, even when paired with food on rare occasions can alter foraging behavior in this species. Individuals could adjust their foraging behavior by tending more or less intensely to specific sensory information. This generalist predator could change major components of its foraging behavior through learning, but the effect of learning on behavior appears mediated by web structure. Learning combined with variation in web structure could explain the substantial individual differences in attack behavior we observe in this species.

## INTRODUCTION

Individuals from many species and populations vary in their diet or behaviour (Bolnick et al., 2003; Réale et al., 2007). These differences are often adaptive and could arise through specialisation (Van Valen, 1965; Roughgarden, 1974; Marchetti & Drent, 2000; Kobler et al., 2009). Specialisation can occur within the lifetime of individuals when they reduce the range of food items consumed or phenotypes expressed to avoid competition with conspecifics (Bergmüller & Taborsky, 2010; Réale et al., 2010) or improve foraging efficiency (Bolnick et al., 2003; Estes et al., 2003; Woo et al., 2008; Araújo et al., 2011; Patrick & Weimerskirch, 2014). This is because the skills required to capture and handle a specific prey item are not always transferable to other prey items (i.e., Darwin’s interference hypothesis; Darwin, 1876; Heinrich et al., 1977; Goulson et al., 1997). Hence, specializing on a narrower set of prey types could reduce foraging costs (Cunningham & Hughes, 1984; Laverty & Plowright, 1988; Goulson et al., 1997; Gegear & Laverty, 1998; Tinker et al., 2009; Dukas, 2019).

How specialization arises is often unclear (Bolnick et al., 2003; Stamps & Groothuis, 2010; Dochtermann et al., 2014; Kok et al., 2019). Past experience can shape an individual’s foraging behavior and diet (e.g., prior success consuming a particular prey type, Persons & Rypstra 2000; Slagsvold & Wiebe, 2007) through learning. Yet, we still have to parse out the role of learning in the emergence of stable foraging specialisation (Werner et al., 1981; Ghalambor et al., 2010; Mery & Burns, 2010; Snell-rood, 2013; Zhang & Hui, 2014; Dukas, 2019; Fraser Franco et al., 2024). One of the most studied forms of learning is associative learning, where the connection between two stimuli and its appropriate behavioral response is reinforced with repeated exposure (Pearce & Bouton, 2001; Dickinson, 2012). Associative learning could lead to specialisation by improving the ability of predators to track the availability of specific prey types in their environment (Mery & Burns, 2010; Snell-rood, 2013; Zhang & Hui, 2014; Dunlap & Stephens, 2016; Dukas, 2019), either by improving the ability of predators to detect or recognize a given type of prey (Hintzman & Block, 1971; Tinker et al., 2009; Dukas, 2019), or by optimizing the sensory cues or spatial locations that predators attend to or their response to such cues (Mery & Burns, 2010). Through these pathways, associative learning could increase foraging success (Werner et al., 1981; Edwards & Jackson, 1994; Holekamp et al., 1997; Morse, 2000; MacDonald, 2007; Reid et al., 2010; Wilson-Rankin, 2015).

A lot of taxa already show a capacity for associative learning (Perry et al., 2013; Álvarez et al., 2017; Ginsburg & Jablonka, 2019; Loy et al., 2021). However, past work has focused on vertebrate cursorial predators, neglecting sedentary predators from taxa thought to have limited cognitive abilities, such as arachnids (Loy et al., 2021). In this study, we assess the ability of Western black widow spiders (*Latrodectus hesperus*) for associative learning in response to vibrational prey cues of various frequencies. Black widow spiders are sedentary predators that spend most of their life on the persistent tridimensional cobweb that they weave (Salomon et al., 2009, 2010). These spiders spend their active time waiting for prey on their web (Kaston, 1970; Uetz, 1992; Hódar & Sánchez-Piñero, 2002; Nelsen et al., 2014). They can detect, identify, and choose whether to attack preys caught in the web using the frequency and strength of vibration (Morley et al., 2016; Vibert et al., 2014, 2016). Spiders adjust the structure of their web and their responsiveness to vibrations according to their hunger level (Zevenbergen et al., 2008; DiRienzo & Montiglio, 2016a). Some spiders, including the Western black widow, already showed their capacity for spatial learning, namely the capacity to associate locations on their webs with the presence of prey (Punzo, 2004; Nakata, 2013; Sergi et al., 2022). Here, we conducted two conditioning experiments where we assessed the ability of black widow spiders to associate vibrations with specific frequencies or applied at specific locations on the web to the presence of a prey. We assess how spiders optimize their response to vibration frequency and location by analysing potential changes in web structure as a result of learning. We hypothesized that individuals would adjust the structure of their webs and their responsiveness towards these frequencies in response to these associations.

## METHODS

### Study system and husbandry

The western black widow spider is found in North America from British Columbia to Mexico and from the west coast to Montana, Wyoming, Colorado and Oklahoma (Miles et al., 2018; Schraft et al., 2021). It is a solitary generalist predator, that mainly feeds on ants (Mackay, 1982), crickets, cockroaches, isopods, and spiders (Salomon, 2011; Trubl et al., 2012). Black widow spider webs are tridimensional and composed of a refuge attached to a sheet of silk threads that is anchored to the substrate by vertical structural threads (Kaston, 1970; Salomon et al., 2010; Vetter et al., 2012). The structural threads are thought to act as defenses against predator and serve to anchor the sheet of silk (Blackledge & Zevenbergen, 2007). The web also includes sticky trap threads between the sheet and the substrate that serve to capture rampant prey (Griswold et al., 1998; Blackledge et al., 2005; Argintean et al., 2006). Individuals further adjust the structure of their web to their hunger level (Heiling & Herberstein, 2000; Zevenbergen et al., 2008) but differ consistently over the number of sticky trap and structural threads that they produce (DiRienzo & Montiglio, 2016b). The number of gum-footed lines has been shown to affect the probability of attacks towards vibrations. Indeed, a web swap experiment testing the attack behavior of individuals on different webs showed that 25 % of the attack probability of the spider could be attributed to the characteristics of their web, but also that individuals differ substantially in their tendency to attack vibrational cues even when assessed over several months (Montiglio & DiRienzo, 2016). Vibrations transmitted by preys on the web range from 0 to 250 Hz, with a mean dominant frequency between 20 and 30 Hz (Vibert et al., 2014). Black widow spiders, like many species, have the capacity to track the distance and direction they travel (Sergi et al., 2021). This means they can later retrace their steps and even take shortcuts or detours while estimating their relative position, thereby providing the bases for spatial learning.

We used western black widow spiders collected in Davis, California. We housed the adult female spiders in 7.7 x 11.7 cm (540 m) individual plastic containers, including a triangular black cardboard refuge, at 21-26°C and 8-53 % humidity under a (12h:12h) photoperiod (Blackledge & Zevenbergen, 2007; DiRienzo & Montiglio, 2016a; DiRienzo & Montiglio, 2016b). We fed the spiders with 1 adult cricket (*Gryllodes sigillatus*) every other week before the start of the experiment. We ascertained that all subjects were sexually mature by inspecting their epigyne visually. At the end of the experiment, we anaesthetized the spiders with CO_2_ before weighting them to the nearest 0.1 mg with an analytical balance (Ohaus Pioneer Precision) and photographing their right front leg on graduated paper. We later used ImageJ to measure the length of their leg as a measure of body size (Schneider et al., 2012).

### Overview of experiments

We performed two experiments where we assessed the effect of repeated conditioning events on the tendency of spiders to attack cues of different frequencies. A first experiment exposed a group of individuals to vibrational cues with specific frequencies twice per week for 8 weeks (yielding 16 conditioning events). Within this experiment, we paired a 250Hz vibrational cue with a cricket at a standardized location on the web for an experimental group. We also paired the 250Hz vibrational cue with a cricket but applied the cue in another location on the web, equidistant to the first one from the refuge for another experimental group. Lastly, for a third experimental group, we randomly paired a different vibrational cue of either 25, 100, 175 or 250Hz and one of the two positions on the web with each presentation of a cricket each week thereby preventing any association between frequency or location with the presence of a prey on the web. The third group acted as control and should have prevented the individuals from associating a specific location or frequency with the presence of food. This first experiment tested whether individuals could learn to associate a specific frequency (250Hz) with the presence of a prey. However, it exposed all individuals to novel frequencies ranging from 25Hz to 250Hz. Thus, all individuals could learn to recognize these frequencies and potentially to associate higher frequencies with the occasional presence of food, irrespective of the treatment. Therefore, to complement this first experiment, we ran a second one where we exposed another group of individuals to two frequencies only (25Hz and 250Hz). In half of the subjects, we always paired the 250Hz with a cricket. For the other half, we paired the cricket with the 25Hz. This experiment was designed to determine whether it was easier to associate some frequencies (25Hz) with food than others. In nature, prey generate vibrations around 25Hz (Vibert et al., 2014). Thus, webs or sensory biases might facilitate learning for cues at 25Hz compared to 250Hz.

We followed the same general procedure for each experiment. We weighted the spiders and placed them on standardized cardboard frame to allow them to spin a web for seven days. We then assessed web structure (see below “web assays”). While the spiders were on their webs, we scored their behavior in response to a range of 4 frequencies (25Hz, 100Hz, 175Hz, and 250Hz) and 2 locations on the web (for experiment 1 only, see Figure 1, yielding 8 different tests for experiment 1 and 4 different tests for experiment 2). We repeated these behavioral assays multiple days on each individual (see “behavioral assays” below). After this initial round of testing, we allocated the spiders randomly to one of three treatments (n=20 per treatment) in experiment 1 or to one of the two treatments (n = 31 per treatment) in experiment 2 and conducted the treatments for 8 weeks twice a week. Each spider thus had 16 learning opportunities. During these learning opportunities, we anesthetized the crickets, placed them on specific location on the web and applied the vibrational cue directly on their body. We then left them on the webs for 24h. Most of the spiders would either attack the crickets right away or would investigate a bit later and were seen eating. After the conditioning treatment, we placed each spider on a new web rack to let it produce a new web. We assessed web structure and attack behavior as above. Lastly, we measured spiders and weighted them before placing them back in their regular housing.

**Figure 1.**
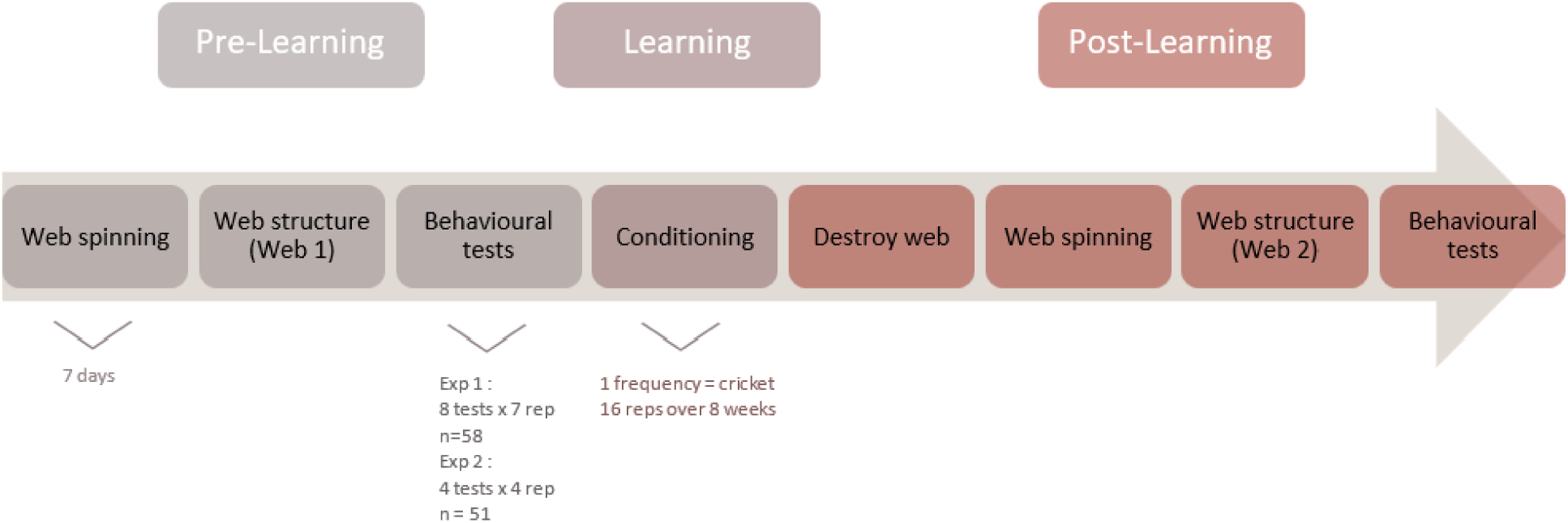
Sequence and experimental design for two experiments on black widow spiders. Subjects underwent the same behavioural tests and web measurements before and after a conditioning phase of 8 weeks. In both experiments, spiders spun webs for 7 days before web measurements and behavioural tests.

### Web assays

Spiders spun their web for 7 days in a standardized cardboard frame (31 cm length × 17 cm height × 24 cm width), the bottom of which was grided (3.8 cm^2^ i.e. 1.5 in). Previous work showed that web structure and the magnitude of individual differences are stable after 7 days (Toupin et al., 2022) while minimizing changes in web structure arising from the drag lines left by spiders as they move on their web, or the damages to the webs by prey and their repairs by individuals (Benjamin & Zschokke, 2002; Eberhard et al., 2008). We assessed web structure by counting the number of trap threads anchored at the bottom of the frame (Blackledge & Zevenbergen, 2007; Zevenbergen et al., 2008; DiRienzo & Montiglio, 2016a; DiRienzo & Montiglio, 2016b; Montiglio & DiRienzo, 2016).

### Behavioral assays

For both experiments, we tested individuals repeatedly for their attack behavior while on their webs before and after the conditioning treatments. We determined the order in which we tested individuals and presented the tests randomly. We created the vibrational cues using the Audacity program (Audacity Team, 2022). Cues consisted of 3 3-second bursts of a given frequency (25, 100, 175 and 250 Hz) with each burst increasing in amplitude. We spaced bursts with 1 sec of silence. These frequencies were within the range of vibrations that spiders encounter in their natural habitat (Vibert et al., 2014). Previous studies showed that similar cues led to attack behavior reliably in black widow spiders, with individuals exhibiting substantial consistent differences in their average attack probability (DiRienzo & Montiglio, 2016a; DiRienzo & Montiglio, 2016b; Montiglio & DiRienzo, 2016; DiRienzo et al., 2020).

We applied the vibrational cues to the web via a metal rod attached to a small speaker (HP DHS-2111; see also (Vibert et al., 2014, 2016). We applied the signal on a single thread that was the closest to predetermined positions on the web. Using the grided bottom of the web rack, we scored the attack behavior of each spider based the presence of a reaction and on the distance traveled from their refuge towards the signal during its duration (Table 1). We waited for a minimum of 10 seconds until the spider was back in her refuge before conducting the next test. We randomized the order in which we tested all individuals each day and randomized the order in which we exposed each individual to the different frequencies. In experiment 1, we tested all spiders daily for 4 and 3 consecutive days separated by a one-day break, for each of the 4 frequencies and 2 locations, thus yielding 8 different tests for each individual and temporal phase (before and after the conditioning treatments). Because preliminary analyses of experiment 1 showed that the 4 consecutive days yielded enough information on individual hunting behavior, we tested all spiders daily for 4 consecutive days for each of the 4 frequencies in experiment 2. In this experiment, we applied all cues at the same location on the web, yielding 4 different tests for each replicate.

**Table 1.**
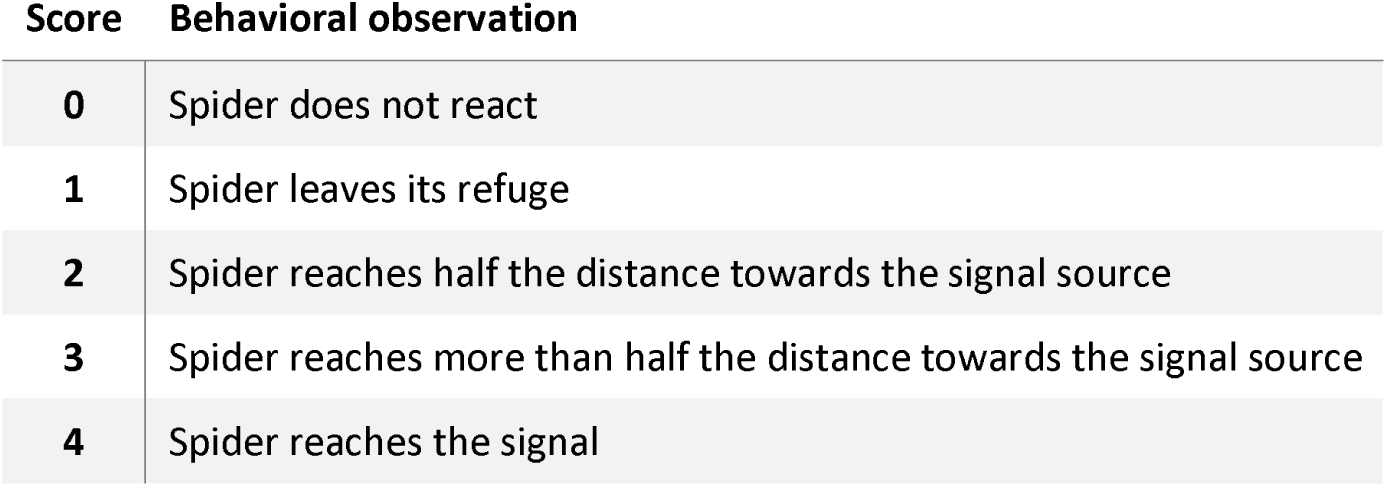
Ethogram quantifying attack behavior in Western black widow spiders towards a vibrational cue (see also Toupin et al., 2022, Ratz et al., 2025).

| Score | Behavioral observation |
| --- | --- |
| 0 | Spider does not react |
| 1 | Spider leaves its refuge |
| 2 | Spider reaches half the distance towards the signal source |
| 3 | Spider reaches more than half the distance towards the signal source |
| 4 | Spider reaches the signal |

### Statistical analyses

We first assessed individual differences in responses to vibrational cues before any experimental treatment. We pooled behavioral assays from both experiments conducted before any experimental treatment into a single generalized linear mixed model analysing individual behavioral scores (ranging from 0 to 4) as a function of the frequency of the vibrational cue (25, 100, 175, or 250Hz, modelled as a categorial variable), the time of day (the number of minutes since midnight, modeled as a continuous variable standardized to a mean of 0 and a standard deviation of 1), and the experiment (1 or 2, binary variable). We included the individual spider identifier, the test date, and an observation-level categorical variable as random effects on the intercept to account for individual differences, temporal variability, and over-dispersion respectively. We modelled behavioral scores using a binomial error distribution (considering the number of successes out of 4 trials). Preliminary work has showed that analysing these scores, the attack probability (i.e., whether the spider scored a 4 or not), or reaction probability (i.e., whether the spider scored 0 or 1 and more) yields qualitatively similar results. To determine whether individuals varied consistently in their response toward the different frequencies, we compared this model with one including random slopes. That is, the effect of the frequency on attack behavior was allowed to vary from one individual to the other. We compared the fit of the two models (with and without random slopes) using a likelihood ratio test. We also computed the repeatability of individual differences in attack behavior on the latent scale following the formula provided for binomial distributions with additive dispersion parameters in Nakagawa and Schielzeth (2010).

Second, we assessed the effect of our experimental treatments in experiment 1 on the number of trap threads in spider webs. We modelled this variable with a Poisson error distribution, as a function of the 6 experimental groups obtained by crossing the temporal phase (before and after conditioning), the reward treatment (control or rewarded at 250Hz), and the location where the stimulus was applied (left or right of the refuge). We also included individual and observation identifiers as random effects to account for pseudo replication and overdispersion respectively. We also used all behavioral assays conducted with a frequency of 25Hz and 250Hz from experiment 1 to assess the effect of conditioning on behavioral scores. We analysed the behavioral scores using a generalized mixed model as above, as a function of a categorical variable identifying experimental groups of observations. We constructed this variable by crossing the experimental treatment (control, left-trained, or right-trained), the temporal phase (before or after the treatments) and frequencies tested (25Hz or 250Hz), thus yielding a 12 different levels (observations before treatment in control individuals at 25Hz, before treatment in control individuals at 250Hz, before treatment in left-trained individuals at 25Hz, etc.). We then constructed 11 orthogonal planned contrasts to assess the differences in behavioral scores among observations made a) at 25Hz vs 250Hz (1 contrast); b) before vs after the treatments at each frequency (2 contrasts); c) on control and trained individuals for each cue frequency and temporal phase (before and after at 25Hz and 250Hz, 4 contrasts); and d) on left-trained and right-trained individuals for each cue frequency and temporal phase (4 contrasts). In this dataset, spiders were tested at two locations on their webs, left or right of the refuge, so we included location as a binary variable in our model and tested for a two-way interaction between position and our categorical variable identifying experimental groups. This interaction allowed us to assess the effect of spatial learning treatment on spider attack behavior. Finally, we included the number of trap lines on the webs during the assays as a standardized continuous variable to account for the impact of web structure on behavioral scores. We estimated the repeatability of random effects as above, following Nakagawa and Schielzeth (2010).

Third, we analysed observations from experiment 2 in a similar way as experiment 1. We analysed the impact of the conditioning treatments (25 or 250Hz) on the number of trap threads in webs as a Poisson process. The model included the experimental groups as a categorical fixed effect, and individual and observation identifiers as random effects. We also analysed the behavioral scores of spiders during the behavioral assays towards 25Hz and 250Hz stimuli as a function of a categorical variable identifying experiment groups of observations as above. We constructed this variable by crossing the experimental group (25Hz or 250Hz-trained individuals), the temporal phase (before or after treatments) and cue frequency (25Hz or 250Hz). We then specified 7 planned contrasts to assess the difference in scores among observations made a) at 25Hz or 250Hz (1 contrast); b) before or after the experimental treatments for each cue frequency (2 contrasts); and c) among the two experimental groups (25Hz and 250Hz-trained individuals) for all phases and cue frequencies (4 contrasts). All observations were made at a single location on the web (right of the refuge) in this experiment. We also included the number of trap threads as a continuous standardized covariable in the model. We calculated the repeatability of random effects as above.

In all models, we tested the statistical significance of random effects using likelihood ratio chi^2^ tests. We ran all models in R 4.3.1 (R Development Core team, 2023) using the lme4 library (Bates et al., 2015). All models initially included the order in which we tested individuals and the order in which we tested frequencies for each individual. However, since these had no effect on the scores, we removed these variables from all analyses.

### Ethical approval

Work presented in this study was not reviewed by an institutional comity because it concerns invertebrate organisms only. This work involved a minor modification of the regular feeding routine of all subjects in captivity. Therefore, it did not have any welfare or environmental implications. This research adheres to the ASAB/ABS Guidelines for the ethical treatment of nonhuman animals in behavioural research and teaching (https://doi.org/10.1016/S0003-3472(24)00376-2) and the ARRIVE Guidelines (https://arriveguidelines.org/arrive-guidelines).

## RESULTS

### Individual differences in responses towards vibrational cues

The model including individual random slopes did not provide a better fit than the model with random intercepts only (Log-likelihood ratio = 5.41, d.f. = 9, p = 0.796). Thus, spiders differed consistently in their overall tendency to respond to cues, but all individuals responded similarly to the different frequencies (see Table S1 for the full model). Repeatability of individual differences was 33.86 % on the latent scale.

### Experiment 1: Effects of spatial and frequency conditioning on web structure and attack behavior

The spiders produced webs with fewer trap threads after the conditioning compared to the beginning of the experiment (estimate = −1.28 ± 0.21, z = −6.03, p < 0.001; see Table S2 and figure 2a). Beyond this temporal effect, the conditioning treatment did not affect the number of trap threads present on the webs.

**Figure 2.**
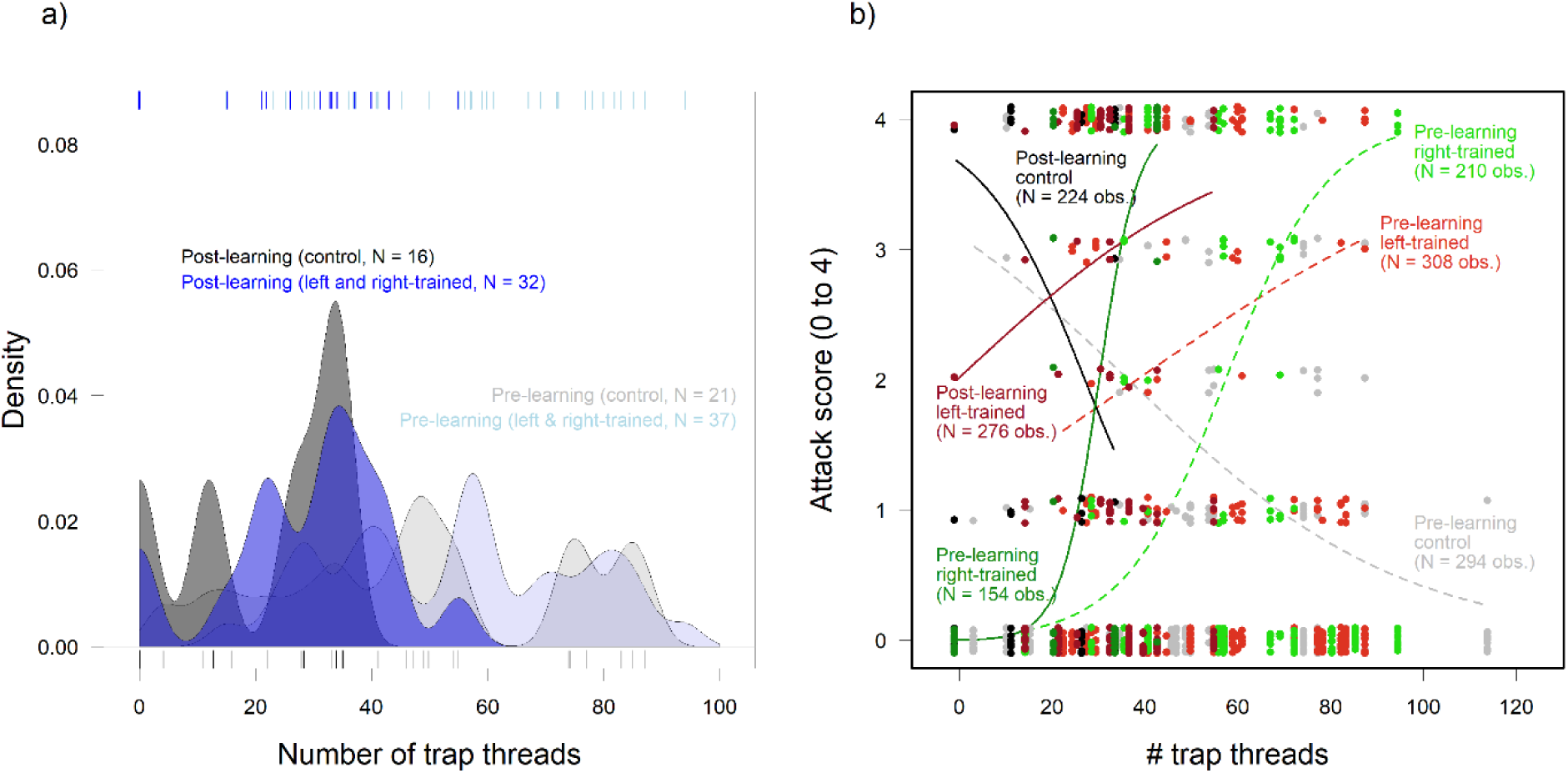
a) Individuals in the control (gray) and experiment (blue) groups decreased the number of trap threads after the conditioning treatment (higher saturation colors) compared to the beginning of the experiment (lower saturation colors) irrespective of their treatment; b) Conditioning in spiders from the experimental groups (left-trained, in red; right-trained, in green) increased the effect of the number of trap threads on their responsiveness during the behavioral assays. Spiders in the control group (control, gray/black) did not show a change in the effect of the number of trap threads on their responsiveness.

The two-way interactions between the experimental group categorical variable and spatial location of cues on the webs was weak and non-significant (Chi^2^= 4.71, d.f. = 11, p = 0.944). We thus removed it from the final model. Unexpectedly, the inspection of residuals led us to detect a strong trend for a two-way interaction between experimental group variable and the number of trap threads. Therefore, we included this interaction in our final model (Chi^2^ test = 18.67, d.f. = 11, p = 0.067; Table 2).

**Table 2.**
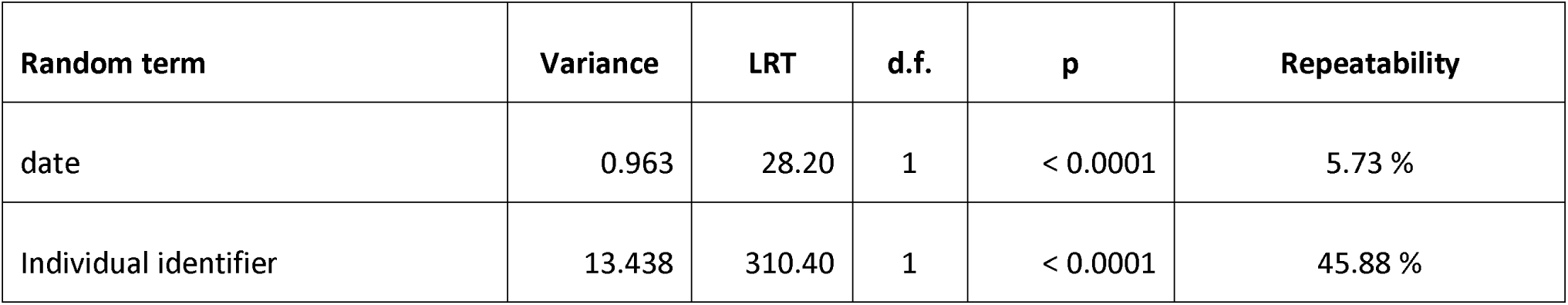
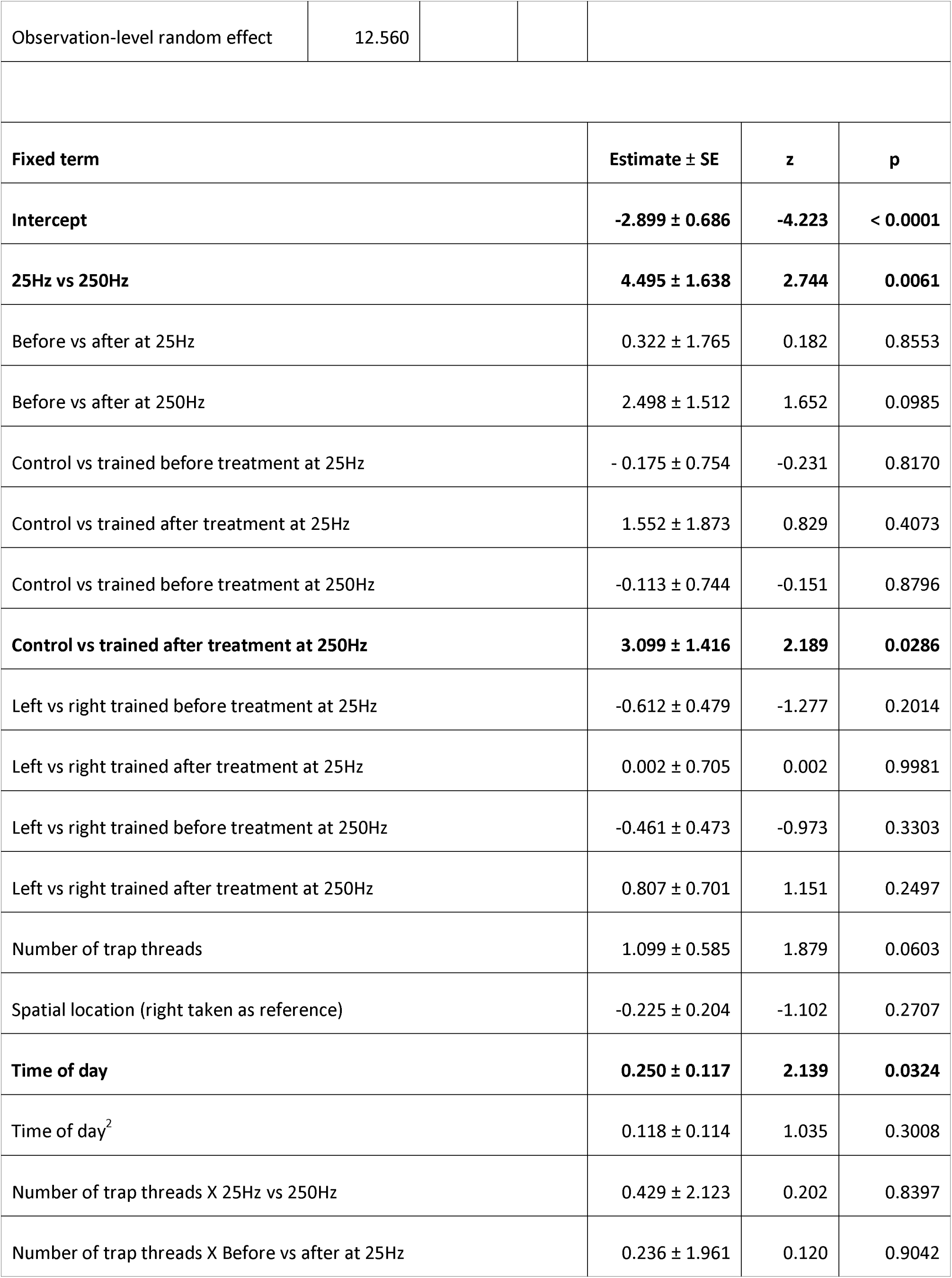
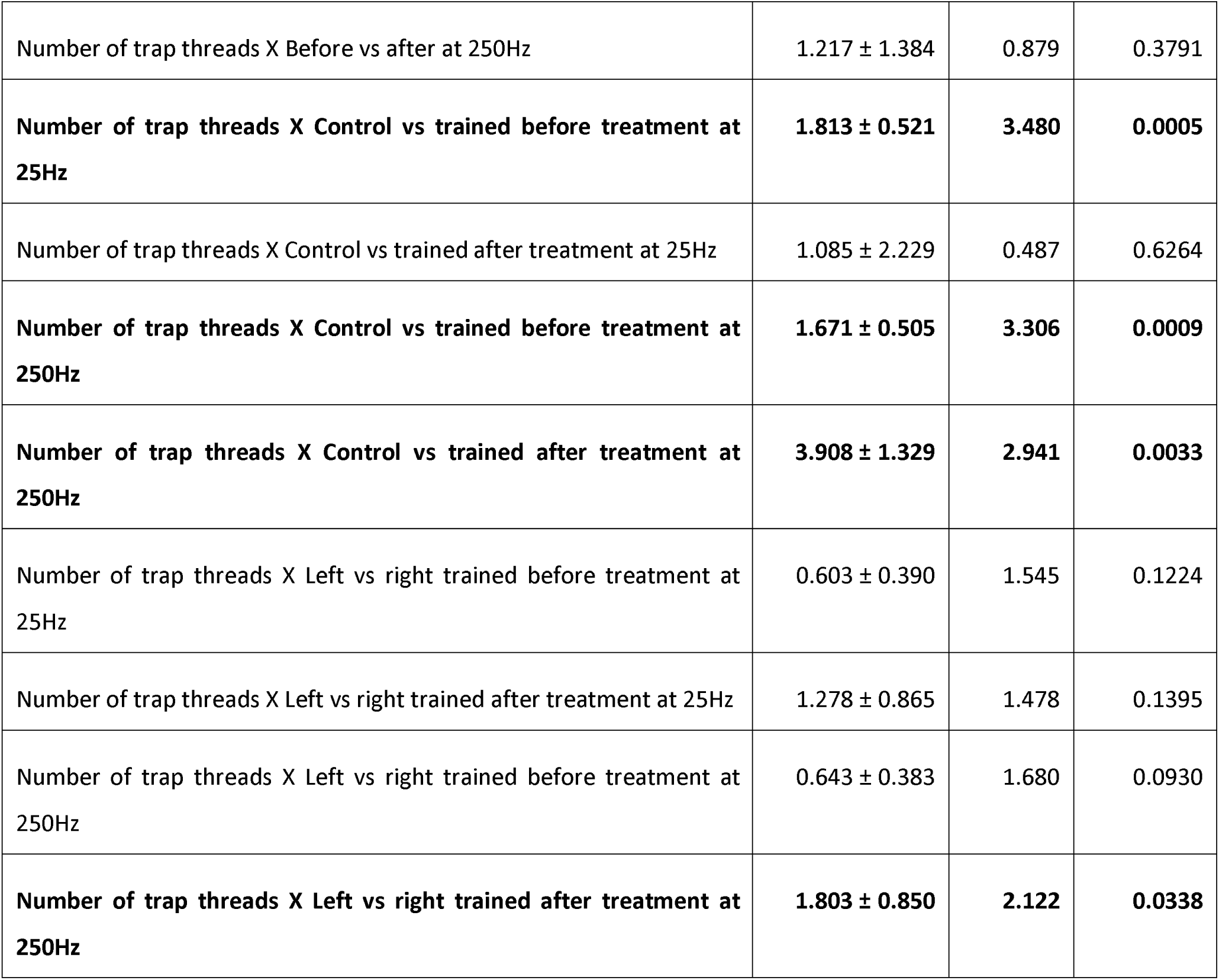
Generalized mixed-effect model analyzing the effect of spatial and 250Hz frequency conditioning on the responsiveness of black widow spiders towards vibrational cues in experiment 1 (2224 observations on 58 individuals on 14 different days). Conditioning treatment for the experimental group represents the effect of receiving a prey reward associated with a frequency of 250 Hz 16 times in one location of the web.

| Random term | Variance | LRT | d.f. | p | Repeatability |
| --- | --- | --- | --- | --- | --- |
| date | 0.963 | 28.20 | 1 | < 0.0001 | 5.73 % |
| Individual identifier | 13.438 | 310.40 | 1 | < 0.0001 | 45.88 % |

|  |  |  |  |
| --- | --- | --- | --- |
| Observation-level random effect | 12.560 |  |  |
| **Fixed term** | **Estimate ± SE** | **z** | **p** |
| **Intercept** | **-2.899 ± 0.686** | **-4.223** | **< 0.0001** |
| **25Hz vs 250Hz** | **4.495 ± 1.638** | **2.744** | **0.0061** |
| Before vs after at 25Hz | 0.322 ± 1.765 | 0.182 | 0.8553 |
| Before vs after at 250Hz | 2.498 ± 1.512 | 1.652 | 0.0985 |
| Control vs trained before treatment at 25Hz | - 0.175 ± 0.754 | -0.231 | 0.8170 |
| Control vs trained after treatment at 25Hz | 1.552 ± 1.873 | 0.829 | 0.4073 |
| Control vs trained before treatment at 250Hz | -0.113 ± 0.744 | -0.151 | 0.8796 |
| **Control vs trained after treatment at 250Hz** | **3.099 ± 1.416** | **2.189** | **0.0286** |
| Left vs right trained before treatment at 25Hz | -0.612 ± 0.479 | -1.277 | 0.2014 |
| Left vs right trained after treatment at 25Hz | 0.002 ± 0.705 | 0.002 | 0.9981 |
| Left vs right trained before treatment at 250Hz | -0.461 ± 0.473 | -0.973 | 0.3303 |
| Left vs right trained after treatment at 250Hz | 0.807 ± 0.701 | 1.151 | 0.2497 |
| Number of trap threads | 1.099 ± 0.585 | 1.879 | 0.0603 |
| Spatial location (right taken as reference) | -0.225 ± 0.204 | -1.102 | 0.2707 |
| **Time of day** | **0.250 ± 0.117** | **2.139** | **0.0324** |
| Time of day<sup>2</sup> | 0.118 ± 0.114 | 1.035 | 0.3008 |
| Number of trap threads X 25Hz vs 250Hz | 0.429 ± 2.123 | 0.202 | 0.8397 |
| Number of trap threads X Before vs after at 25Hz | 0.236 ± 1.961 | 0.120 | 0.9042 |

|  |  |  |  |
| --- | --- | --- | --- |
| Number of trap threads X Before vs after at 250Hz | 1.217 ± 1.384 | 0.879 | 0.3791 |
| **Number of trap threads X Control vs trained before treatment at 25Hz** | **1.813 ± 0.521** | **3.480** | **0.0005** |
| Number of trap threads X Control vs trained after treatment at 25Hz | 1.085 ± 2.229 | 0.487 | 0.6264 |
| **Number of trap threads X Control vs trained before treatment at 250Hz** | **1.671 ± 0.505** | **3.306** | **0.0009** |
| **Number of trap threads X Control vs trained after treatment at 250Hz** | **3.908 ± 1.329** | **2.941** | **0.0033** |
| Number of trap threads X Left vs right trained before treatment at 25Hz | 0.603 ± 0.390 | 1.545 | 0.1224 |
| Number of trap threads X Left vs right trained after treatment at 25Hz | 1.278 ± 0.865 | 1.478 | 0.1395 |
| Number of trap threads X Left vs right trained before treatment at 250Hz | 0.643 ± 0.383 | 1.680 | 0.0930 |
| **Number of trap threads X Left vs right trained after treatment at 250Hz** | **1.803 ± 0.850** | **2.122** | **0.0338** |

When tested on a web with an average number of trap threads (i.e. when the scaled number of trap threads was held at 0), the model indicated that spiders mounted stronger responses towards cues at 250Hz compared to 25Hz across all treatments and for both temporal phases (estimate = −2.90 ± 0.69, z = −4.22, p < 0.001, Table 2). Still at average numbers of trap threads, responses did not differ between control and trained individuals at 25Hz neither before (estimate = −0.17 ± 0.75, z = −0.23, p = 0.817) nor after (estimate = 1.55 ± 1.87, z = 0.83, p = 0.407). At 250Hz, we also failed to detect any differences among control and trained individuals before the experiment (estimate = −0.11 ± 0.74, z = −0.15, p = 0.88). However, trained individuals mounted stronger responses toward 250 Hz vibrational cues after treatment compared to the controls (estimate = 3.10 ± 1.42, z = 2.19, p = 0.029).

The two-way interactions between the experimental group categorical variable and the number of trap threads suggested that the conditioning also affected the effect of trap threads on individual responses towards vibrational cues (Figure 2b, Table 2). More trap threads on the webs increased the responses of trained individuals to a greater extent than control individuals at 25Hz and 250Hz before treatment (estimate at 25Hz = 1.81 ± 0.52, z = 3.48, p = 0.0005; estimate at 250 Hz = 1.67 ± 0.51, z = 3.31, p = 0.0009), and this effect was even stronger after the treatment (estimate = 3.91 ± 1.33, z = 2.94, p = 0.003). Interestingly, the number of trap threads had a stronger effect on the responses of individuals to vibrational cues when these were applied right of the refuge (estimate = 1.80 ± 0.85, z = 2.12, p = 0.034).

### Experiment 2: Effect of conditioning on attack behavior at two frequencies

The spiders produced webs with fewer trap threads after the conditioning compared to the beginning of the experiment (estimate = −0.67 ± 0.32, z = −2.10, p = 0.036; see Table S3 and Figure 3a). Before the conditioning treatment, spiders rewarded at 250Hz also produced more trap threads than spiders rewarded at 25Hz (estimate = −0.48 ± 0.22, z = −2.20, p = 0.028). However, this difference disappeared after the conditioning treatment (estimate = 0.22 ± 0.26, z = 0.86, p = 0.390).

**Figure 3.**
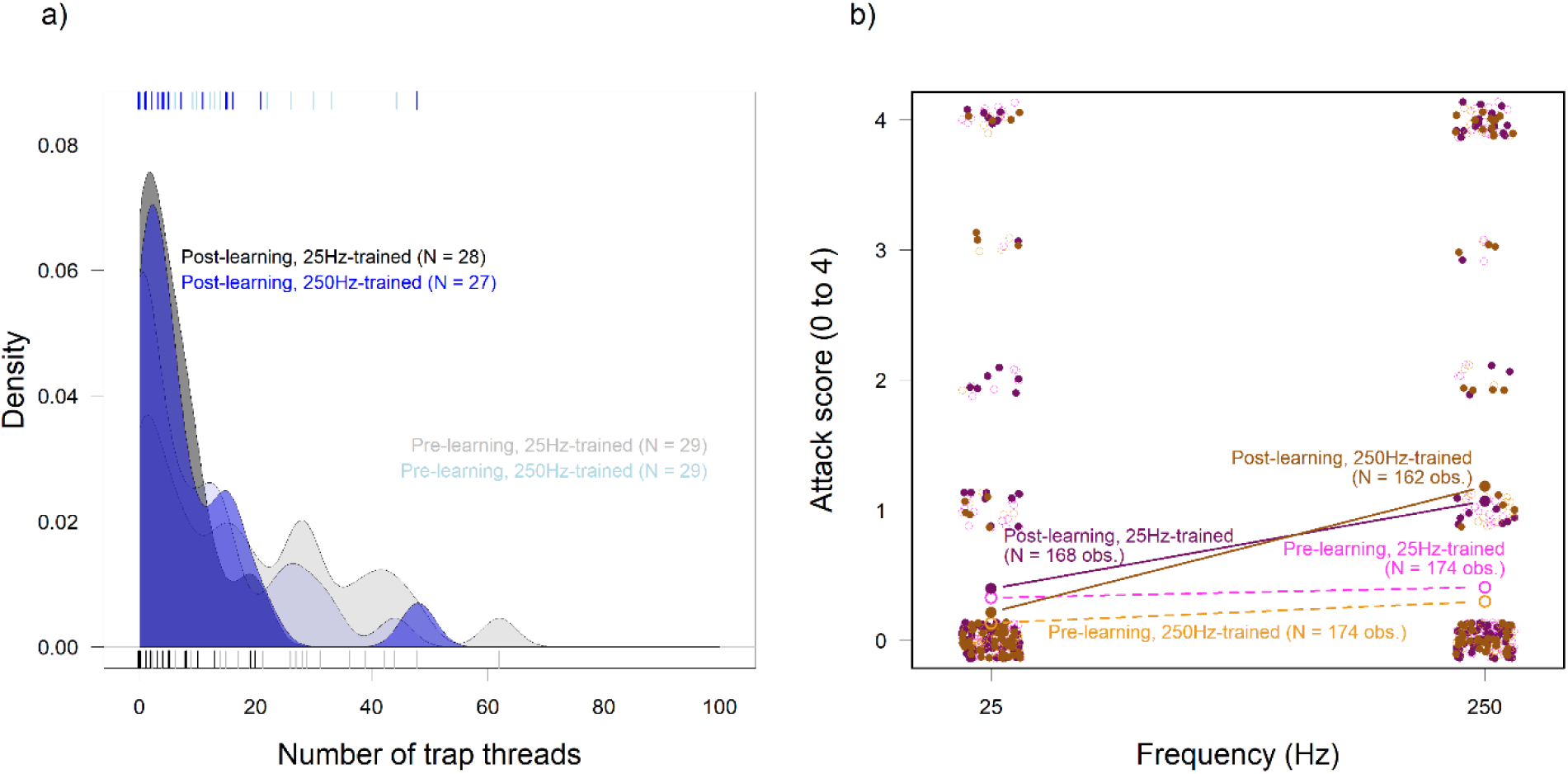
a) Spiders from all groups (conditioning at 25Hz, in gray; conditioning at 250Hz, in blue) decreased the number of trap threads before (lower saturation) and after (higher saturation) the experiment 2. b) Spiders rewarded at 25Hz (pink) or 250Hz (brown) both increased their responsiveness towards 250Hz vibrational cues after conditioning (filled points and solid lines) compared to the beginning of the experiment (empty points and dashed lines).

In a similar way as experiment 1, the number of trap threads appeared to interact with experimental treatments to affect the response of individuals to vibrational stimuli (Chi^2^ = 11.54, d.f. = 7, p = 0.073). However, none of the estimates in the model reached significance (Table S4). For parsimony, we thus excluded this interaction from our model. Analysis of this simpler model showed that individuals responded more readily towards cues at 250Hz than 25Hz (estimate = 2.15 ± 0.78, z = 2.74, p = 0.0061; Table 3; Figure 3b), and this was mostly attributable to individuals increasing their responsiveness toward cues at 250 Hz as a result of the conditioning treatment, irrespective of their experimental group (estimate = 1.40 ± 0.58, z = 2.394, p = 0.0166, Figure 3b). We did not detect any other effect of conditioning treatment on responsiveness. Importantly, spiders rewarded at 25Hz did not increase their responsiveness towards this frequency as a result of the conditioning (estimate = −0.33 ± 0.54, z = −0.60, p = 0.5467) nor did the spiders rewarded at 250Hz increase their responsiveness towards this frequency (estimate = 0.070 ± 0.50, z = 0.14, p = 0.888).

**Table 3.**
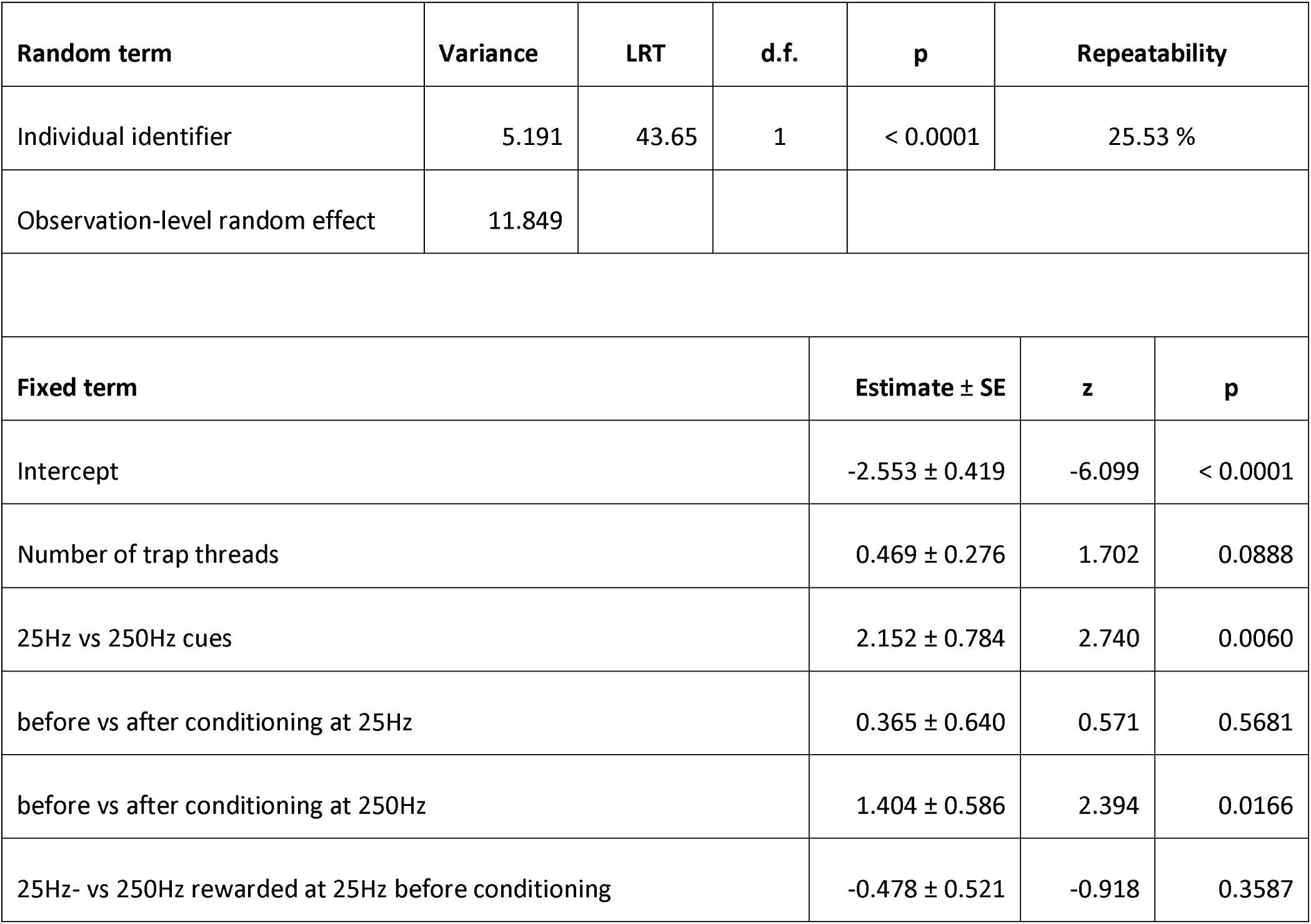
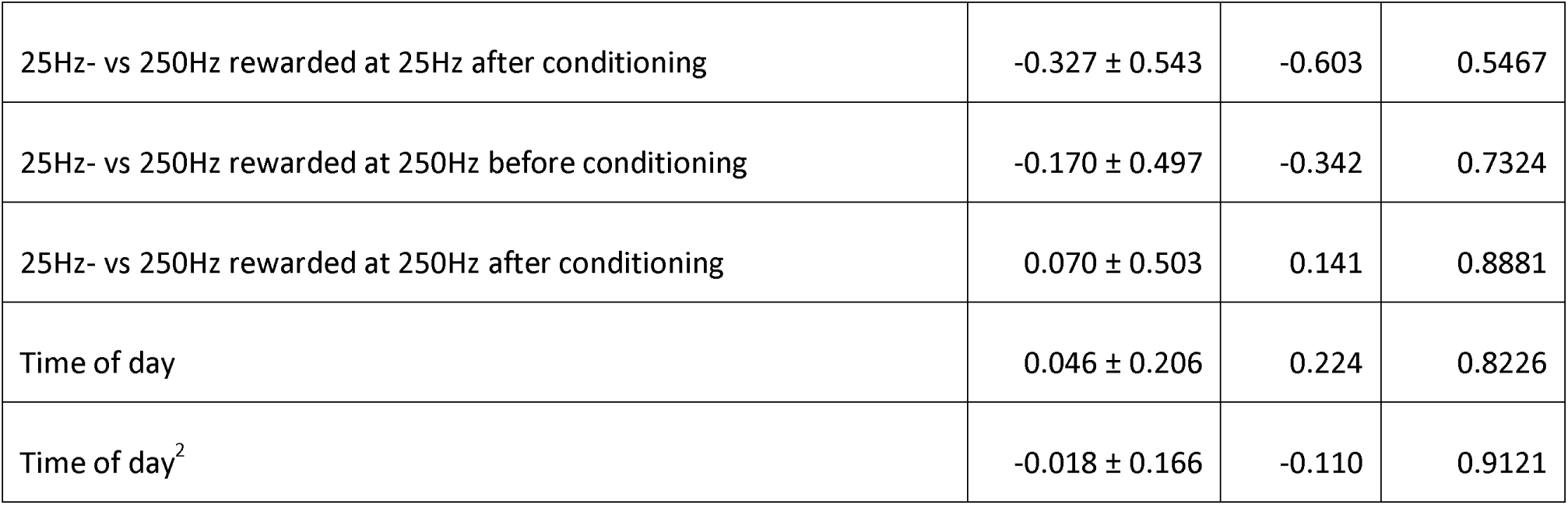
Generalized mixed-effect model analyzing the effect of conditioning at 25Hz or 250Hz on the responsiveness of black widow spiders in experiment 2 (binomial error distribution; 564 observations on 51 individuals). Conditioning treatment represents the effect of receiving a prey reward associated with a frequency 16 times. A date random effect did not explain any variance and was therefore removed to facilitate convergence.

| Random term | Variance | LRT | d.f. | p | Repeatability |
| --- | --- | --- | --- | --- | --- |
| Individual identifier | 5.191 | 43.65 | 1 | < 0.0001 | 25.53 % |
| Observation-level random effect | 11.849 |  |  |  |  |
| Fixed term | Estimate ± SE | | z | p | |
| Intercept | $-2.553 \pm 0.419$ | | -6.099 | < 0.0001 | |
| Number of trap threads | $0.469 \pm 0.276$ | | 1.702 | 0.0888 | |
| 25Hz vs 250Hz cues | $2.152 \pm 0.784$ | | 2.740 | 0.0060 | |
| before vs after conditioning at 25Hz | $0.365 \pm 0.640$ | | 0.571 | 0.5681 | |
| before vs after conditioning at 250Hz | $1.404 \pm 0.586$ | | 2.394 | 0.0166 | |
| 25Hz- vs 250Hz rewarded at 25Hz before conditioning | $-0.478 \pm 0.521$ | | -0.918 | 0.3587 | |

|  |  |  |  |
| --- | --- | --- | --- |
| 25Hz- vs 250Hz rewarded at 25Hz after conditioning | -0.327 ± 0.543 | -0.603 | 0.5467 |
| 25Hz- vs 250Hz rewarded at 250Hz before conditioning | -0.170 ± 0.497 | -0.342 | 0.7324 |
| 25Hz- vs 250Hz rewarded at 250Hz after conditioning | 0.070 ± 0.503 | 0.141 | 0.8881 |
| Time of day | 0.046 ± 0.206 | 0.224 | 0.8226 |
| Time of day<sup>2</sup> | -0.018 ± 0.166 | -0.110 | 0.9121 |

## DISCUSSION

In this study, we analysed the potential consequences of associative learning on two major components of the foraging behavior in Western black widow spiders: the web structure and propensity to respond to vibrations. The first experiment showed that spiders do not adjust the structure of their web nor their response to specific web locations when we applied prey items and vibrations at specific locations on the web. Instead, all spiders increased their responsiveness to vibrations at 250 Hz after repeated exposures to this frequency irrespective of their experimental treatment. Hence, this experiment suggests that repeated exposures, even when paired with food on rare occasions can alter foraging behavior in this species. Exposing spiders to prey items associated with a 250Hz increased the effect of trap threads on responsiveness itself. This suggest that individuals adjust their foraging behavior by tending more or less intensely to specific sensory information. A second experiment largely replicated our findings. Spiders did not change the structure of their webs but increased their responsiveness towards vibrations at 250Hz, irrespective of their experimental treatment. This second experiment did not demonstrate changes in responsiveness following conditioning at 25Hz. It might be easier for these animals to learn to distinguish or attend to higher frequencies than lower ones.

### Evidence for associative learning in spiders

Exposing Western black widow spiders weekly to 250Hz vibrations for 8 weeks increased their responsiveness toward vibrational cues with this frequency. We witnessed this increase in responsiveness when we paired systematically this frequency with a food item (left and right-trained experimental groups in experiment 1 and 250Hz-trained group in experiment 2), but also when we paired all frequencies with a food reward (control group in experiment 1), and even in absence of any reward (25Hz-trained group in experiment 2). In contrast, we did not observe this increase in responsiveness toward 25Hz. Thus, it seems that repeated exposure to a food item can sensitize individuals but only for higher frequencies. The only study that has recorded vibrations from prey in Western black widow spider webs suggests that prey typically produce vibrations with a dominant frequency between 20 and 30Hz (Vibert et al., 2014). Thus, exposing spiders to 25Hz vibrations could fail to cause any change in their foraging behavior because this is within the frequencies they already associate with prey items. In contrast, 250Hz vibrations might have been too novel a stimulus for the individuals to elicit foraging responses prior to repeated exposure. This hypothesis however, does not account for the higher responsiveness of spiders towards 250Hz compared to 25Hz. In our experiments, all frequencies were applied on the webs in a random order but always sequentially (i.e., in the control group in experiment 1 and all groups in experiment 2). That is, we exposed spiders to all frequencies before moving on to the next subject. Hence, it is possible that spiders associated the 250Hz with a prey item and increased their responses by associating all vibrational cues with the presence of a prey. However, this does not account for the lack of response of subject to repeated exposure to lower frequencies (25Hz).

### Learning through response to web structure

Our experiment yielded an unexpected and novel result. Exposing spiders to vibrations and prey items in tandem altered the effect of web structure on responsiveness towards vibrations. Trap threads are hypothesized to function in prey capture, either by trapping prey in glue droplets, or by providing information on the presence and location of potential prey below the web. Spiders are known to increase the number of trap threads on their webs when hungry (Blackledge & Zevenbergen, 2006), and webs with more trap threads are known to enable higher levels of responsiveness toward vibrations (Montiglio and DiRienzo, 2016). In our first experiment, the number of trap threads had a negative effect on responsiveness at 250Hz prior to the treatment, and this effect became clearly positive after the treatment. Hence, spiders could adjust their foraging behavior adaptively by tending more closely to the vibrational cues provided by trap threads on their web. We observed a similar trend in our second experiment but failed to replicate its statistical significance. Webs are known to play an important role in the sensory ecology of spiders (Witt, 1975; Foelix, 1996), and they are believed to have evolved to allow a greater sensitivity to the vibrations generated by prey or mates (Wignall & Herbertstein, 2013, Vibert et al., 2014). Earlier work has shown that we can condition spiders to respond to or to avoid certain vibratory cues (e.g., cross spiders, Bays, 1962; jumping spiders, Long et al., 2015); wolf spiders, Stoffer et al., 2021). Our results indicate that sedentary spiders can also develop some preference by attacking some preys more frequently than others on their web.

Interestingly, our conditioning treatment, pairing prey items with a 250Hz vibration did not alter the number of trap threads. This suggests that Western black widows can adjust their foraging behavior mainly by tending more closely to some cues born by their webs rather than adjusting the structure of their webs. Our finding is surprising given that many orb weaving spiders alter their web according to the size of prey or capture frequency (Sandoval, 1994; Venner et al., 2000; Tso et al., 2007; Blamires, 2010). Orb weaving spiders renew their web each day and this could enable them to use past experience when rebuilding their webs. In contrast, black widow spiders build persistent cobwebs (Schraft et al. 2021), which might be too costly to alter past a certain level. In our experiments, all spiders decreased the number of trap threads they built in their webs at the end of the experiment compared to its beginning, probably because they became more satiated across the repeated feeding events (Blackledge & Zevenbergen, 2007; DiRienzo & Montiglio, 2016a). In contrast to our results on frequencies, black widow spider did not adjust their responsiveness based on cue location on their webs. Applying prey items and vibrations to the left or right of the refuge did not alter individuals’ responsiveness to these locations on their webs, which is in accordance with previous work (Thompson et al., 2020). Black widow spiders have the capacity for path integration and spatial learning (Sergi et al. 2022). However, such spatial learning could depend on the way cues are applied to the web.

We also documented consistent individual differences in responsiveness during the course of both experiments. Some individuals responded more to vibrations than others. Past work has shown that individuals differed in their tendency to attack a single vibrational cue (DiRienzo and Montiglio 2016b). We extend these results by reporting that these differences in responsiveness are detectable across a range of frequencies. The repeatability, quantifying the importance of such individual variation, was well within the range of reported values for behavioral traits (Bell et al., 2009). We also documented individual differences in web structure despite imposing standardized exposures to prey and vibrations in our experiment. Such individual variation in web structure is now well established (DiRienzo & Montiglio, 2016b; Thompson et al., 2020; Toupin et al., 2022). Individuals could either focus on building webs to forage or to protect themselves (Zevenbergen et al., 2008). These individual differences in responsiveness and web structure could generate individual differences in food niche. In spiders, prey capture specialisation has been shown at the species or population level (Uetz et al., 1978; Olive, 1980; Líznarová et al., 2013). For example, a spider’s legs, fangs, overall size or web characteristic would make them efficient at capturing a specific prey type. An interesting avenue for future research will be to analyse individual specialisation in diet and prey capture (Uetz, 1992; Beleyur et al., 2015).

## CONCLUSION

We reported on two experiments investigating the ability of spiders to adjust their foraging behavior in response to repeated exposure of prey items associated with specific vibration frequencies. We observed that Western black widow spiders do indeed increase their responsiveness toward higher frequencies but that they do not alter their web structure nor respond to prey location. Interestingly, changes in responsiveness toward the vibrations with higher frequencies was mediated by the trap threads built in the webs. Hence, individual spiders could learn to respond preferentially to prey with specific vibrations by adjusting their ability to detect or monitor specific vibrational cues transmitted by trap threads. This study thus suggest that learning plays a complex role in shaping the foraging behavior in this generalist sedentary predator through an interaction with its extended phenotype. Learning combined with variation in web structure could explain the substantial individual differences in attack behavior we observe in this species. In line with a growing body of recent research, our study suggests that learned preferences could explain part of the individual differences in responsiveness in this species and provide the substrate for individual foraging specialization.

## ACKOWLEDGEMENTS

This work was funded by a discovery grant from the National Science and Engineering Research Council (Canada) to POM. The authors thank all current and past members of the Montiglio laboratory for their help caring for the spiders and comments on earlier versions of this manuscript.

## Supplementary material

**Table S1.** Generalized mixed-effect model analyzing the responsiveness of spiders as a function of vibrational cue frequency from experiments 1 and 2. (N = 2236 observations from 109 individuals on 10 days).

| Random term | Variance | LRT | d.f. | p | repeatability |  |
| --- | --- | --- | --- | --- | --- | --- |
| Individual identifier | 7.372 | 388.62 | 1 | < 0.0001 | 33.86 % |  |
| date | 0.464 | 26.44 | 1 | < 0.0001 |  |  |
| Observation-level random effect | 11.106 |  |  |  |  |  |
| Fixed term |  |  |  | Estimate ± SE | z | P |
| Intercept (25Hz taken as reference) |  |  |  | -2.990 ± 1.095 | -2.732 | 0.0063 |
| Frequency 100Hz |  |  |  | 1.175 ± 0.268 | 4.383 | < 0.0001 |
| Frequency 175Hz |  |  |  | 0.984 ± 0.269 | 3.658 | 0.0003 |
| Frequency 250Hz |  |  |  | 0.673 ± 0.272 | 2.478 | 0.0132 |
| Time of day |  |  |  | 0.169 ± 0.114 | 1.487 | 0.1369 |
| Experiment (1 v.s. 2, 1 taken as reference) |  |  |  | -0.116 ± 0.739 | -0.157 | 0.8750 |

**Table S2.** Generalized mixed-effect model analysing the number of trap threads on spider webs as a function of experimental treatments in experiment 1 (Poisson error distribution; N = 80 webs from 58 individuals).

| Random term | Variance | LRT | d.f. | p | repeatability |  |
| --- | --- | --- | --- | --- | --- | --- |
| Individual identifier | 0.181 | 8.088 | 1 | 0.0045 | 49.27 % |  |
| Observation-level random effect | 0.155 |  |  |  |  |  |
| Fixed term |  |  |  | Estimate ± SE | z | P |
| Intercept |  |  |  | 3.441 ± 0.091 | 37.784 | < 0.0001 |
| exp_groups2cont_prevspost |  |  |  | -1.282 ± 0.213 | -6.028 | < 0.0001 |
| exp_groups2cont_pre_TvsExp |  |  |  | 0.105 ± 0.110 | 0.953 | 0.340 |
| exp_groups2cont_post_TvsExp |  |  |  | 0.262 ± 0.207 | 1.264 | 0.206 |
| exp_groups2cont_pre_Exp_gvsd |  |  |  | 0.023 ± 0.067 | 0.337 | 0.736 |
| exp_groups2cont_post_Exp_gvsd |  |  |  | -0.066 ± 0.112 | -0.590 | 0.555 |

**Table S3.** Generalized mixed-effect model analysing the effects of conditioning treatment on the number of trap threads produced by individuals in experiment 2 (Poisson error distribution; N = 101 observations from 58 individuals).

| Random effect | Variance | LRT | d.f. | p | Repeatability |  |
| --- | --- | --- | --- | --- | --- | --- |
| Individual identifier | 0.440 | 0.744 | 1 | 0.388 | 16.66 % |  |
| Observation-level random effect | 1.976 |  |  |  |  |  |
| Fixed term |  |  |  | Estimate ± SE | z | p |
| Intercept |  |  |  | 1.384 ± 0.194 | 7.148 | < 0.0001 |
| Before vs after conditioning |  |  |  | -0.674 ± 0.322 | -2.096 | 0.0361 |
| Before conditioning control group v.s. rewarded group |  |  |  | -0.486 ± 0.221 | -2.198 | 0.0280 |
| After conditioning control group v.s. rewarded group |  |  |  | 0.227 ± 0.264 | 0.859 | 0.3902 |

**Table S4.**
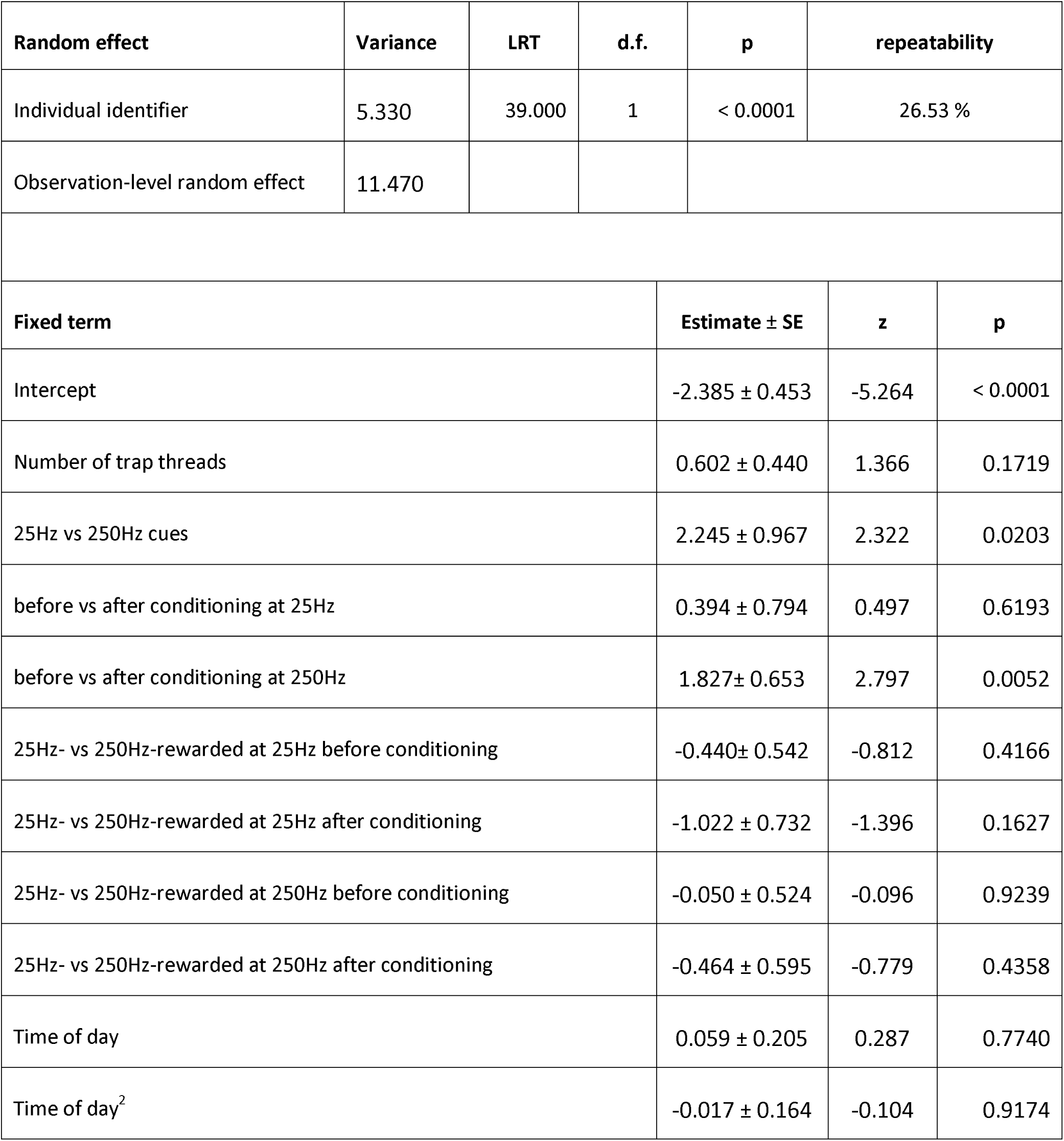
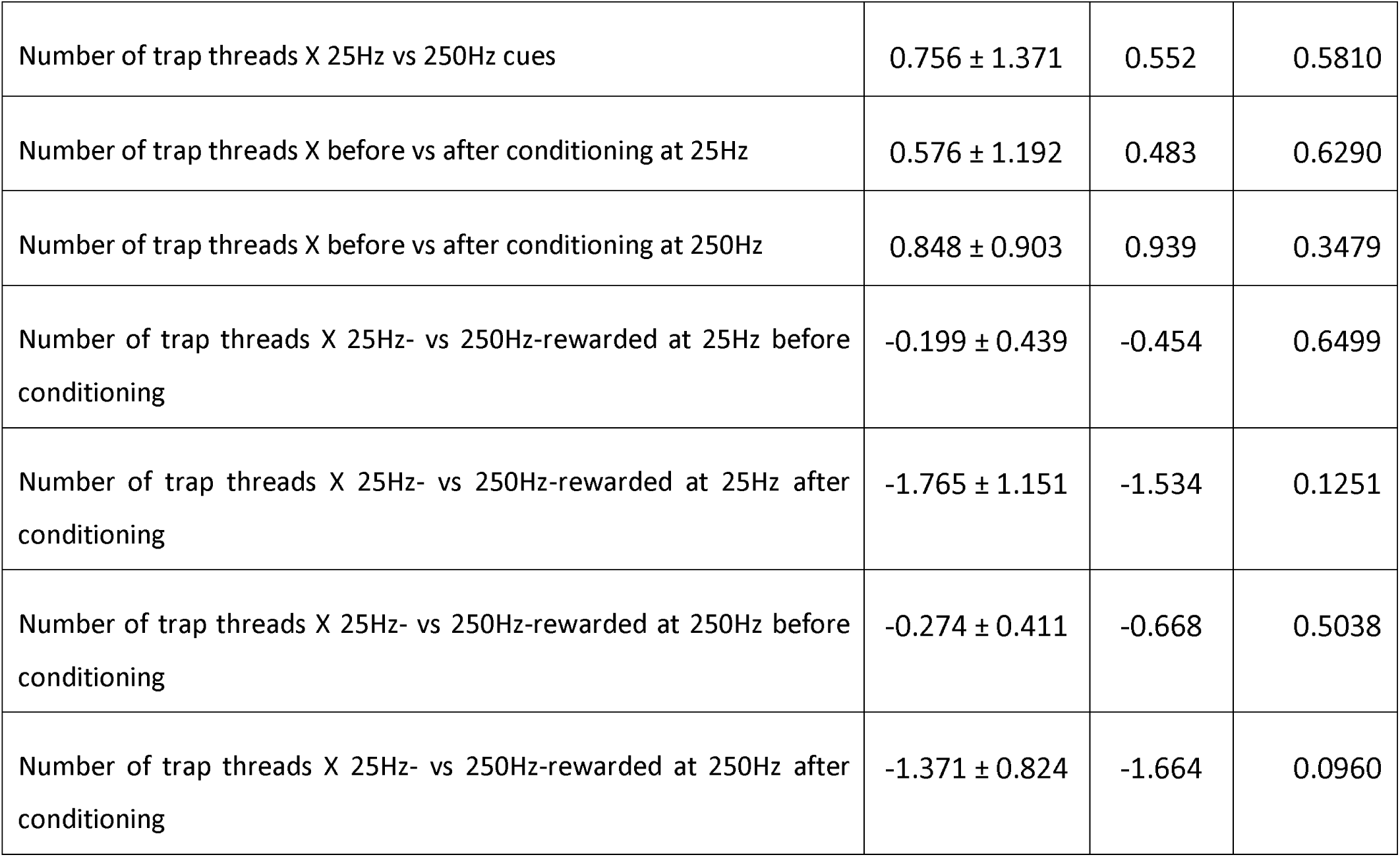
Generalized mixed-effect model analysing the effect of conditioning treatment and the number of trap threads on the responsiveness of individuals toward vibrational prey cues in experiment 2 (binomial error distribution; model including a two-way interaction between the experimental group variable and the number of trap threads; N = 564 observations from 51 individuals).

| Random effect | Variance | LRT | d.f. | p | repeatability |  |
| --- | --- | --- | --- | --- | --- | --- |
| Individual identifier | 5.330 | 39.000 | 1 | < 0.0001 | 26.53 % |  |
| Observation-level random effect | 11.470 |  |  |  |  |  |
| Fixed term |  |  |  | Estimate ± SE | z | p |
| Intercept |  |  |  | -2.385 ± 0.453 | -5.264 | < 0.0001 |
| Number of trap threads |  |  |  | 0.602 ± 0.440 | 1.366 | 0.1719 |
| 25Hz vs 250Hz cues |  |  |  | 2.245 ± 0.967 | 2.322 | 0.0203 |
| before vs after conditioning at 25Hz |  |  |  | 0.394 ± 0.794 | 0.497 | 0.6193 |
| before vs after conditioning at 250Hz |  |  |  | 1.827± 0.653 | 2.797 | 0.0052 |
| 25Hz- vs 250Hz-rewarded at 25Hz before conditioning |  |  |  | -0.440± 0.542 | -0.812 | 0.4166 |
| 25Hz- vs 250Hz-rewarded at 25Hz after conditioning |  |  |  | -1.022 ± 0.732 | -1.396 | 0.1627 |
| 25Hz- vs 250Hz-rewarded at 250Hz before conditioning |  |  |  | -0.050 ± 0.524 | -0.096 | 0.9239 |
| 25Hz- vs 250Hz-rewarded at 250Hz after conditioning |  |  |  | -0.464 ± 0.595 | -0.779 | 0.4358 |
| Time of day |  |  |  | 0.059 ± 0.205 | 0.287 | 0.7740 |
| Time of day<sup>2</sup> |  |  |  | -0.017 ± 0.164 | -0.104 | 0.9174 |

|  |  |  |  |
| --- | --- | --- | --- |
| Number of trap threads X 25Hz vs 250Hz cues | 0.756 ± 1.371 | 0.552 | 0.5810 |
| Number of trap threads X before vs after conditioning at 25Hz | 0.576 ± 1.192 | 0.483 | 0.6290 |
| Number of trap threads X before vs after conditioning at 250Hz | 0.848 ± 0.903 | 0.939 | 0.3479 |
| Number of trap threads X 25Hz- vs 250Hz-rewarded at 25Hz before conditioning | -0.199 ± 0.439 | -0.454 | 0.6499 |
| Number of trap threads X 25Hz- vs 250Hz-rewarded at 25Hz after conditioning | -1.765 ± 1.151 | -1.534 | 0.1251 |
| Number of trap threads X 25Hz- vs 250Hz-rewarded at 250Hz before conditioning | -0.274 ± 0.411 | -0.668 | 0.5038 |
| Number of trap threads X 25Hz- vs 250Hz-rewarded at 250Hz after conditioning | -1.371 ± 0.824 | -1.664 | 0.0960 |

